# Multi-scale analysis of schizophrenia risk loci: Integrating centenarian genomes and spatio-temporal expression profiles suggests the need for adjunctive therapeutic interventions for neuropsychiatric disorders

**DOI:** 10.1101/369090

**Authors:** Chellappa S Anirudh, Ankit Kumar Pathak, Prashant Sinha, Ashwin K. Jainarayanan, Sanjeev Jain, Samir K. Brahmachari

## Abstract

Schizophrenia (SZ) is a debilitating mental illness with multigenic etiology and significant heritability. Despite extensive genetic studies the molecular etiology has remained enigmatic. A recent systems biology study suggested a protein-protein interaction (PPI) network for SZ with 504 novel interactions. The onset of psychiatric disorders is predominantly during adolescence often accompanied by subtle structural abnormalities in multiple regions of the brain. The availability of BrainSpan atlas data allowed us to re-examine the genes present in SZ interactome as a function of space and time. The availability of genomes of healthy centenarians and non-psychiatric ExAC database allowed us to identify the *variants of criticality*. The expression of SZ candidate genes responsible for cognition and disease onset were studied in different brain regions during particular developmental stages. A subset of novel interactors detected in the network was further validated using gene-expression data of post-mortem brains of patients with psychiatric illness. We have narrowed down the list of drug targets proposed by the previous interactome study to 10 proteins. These proteins belonging to 81 biological pathways, are targeted by 34 known FDA approved drugs that have distinct potential for treatment of neuropsychiatric disorders. We also report the possibility of targeting key genes belonging to Celecoxib pharmacodynamics, Gα signaling and cGMP-PKG signaling pathways, that are non-specific to schizophrenia etiology.

## INTRODUCTION

Schizophrenia (SZ) is a complex psychiatric disorder with multi-genic aetiology, affecting almost 1% of the global population (McGrath J *et al*., 2008). It has been clear that the disorder is highly heritable and there is a strong genetic basis, which has been a focus of research over the past decade (Cardno *et al*., 2012). Complex neuropsychiatric disorders like SZ, Major depressive disorder (MDD), Bipolar disorder (BP), Obsessive compulsive disorder (OCD) and Autism spectrum disorder (ASD) are driven by multiple genetic variants across various genomic loci that perhaps interact with environmental factors to produce the disease phenotype (Viswanath *et al*., 2018). The National Human Genome Research Institute (NHGRI) of USA has catalogued 38 genome-wide association studies (GWAS) on SZ (Hindroff *et al*., 2009) revealing the association of common variants with SZ (Girard *et al*., 2012). In addition, the Psychiatric Genomics Consortium (PGC) has identified 108 SZ associated loci (Ripke *et al*., 2014). The molecular mechanisms by which these genetic variations contribute to psychoses could be better understood by studying protein-protein interactions and other molecular interaction networks. Recently, a novel random forest model named High-Confidence Protein-Protein Interaction Prediction (**HiPPIP**) was developed to classify the pairwise features of interacting proteins. The HiPPIP predicted 504 novel PPIs adding to 1397 known PPIs, for 101 SZ candidate genes, presenting a novel theoretical interactome for SZ. A few (pair-wise interactions) were experimentally validated (Ganapathiraju *et al*., 2016). The analysis illustrates that despite the divergent findings of different studies on SZ, a common thread emerges as the genes lead to pathways, through the interaction network. Several genes present in key pathways deduced from the interactome are targets of existing drugs used to manage various chronic diseases (Ganapathiraju *et al*., 2016).

While tissue specific gene expression data from the Stanford Microarray Database (SMD) and Tissue-specific Gene Expression and Regulation (TiGER) database were included to build the HiPPIP model, it still lacked a spatio-temporal information. SZ, a developmental disorder of largely adolescent onset, is associated with subtle structural abnormalities and molecular differences in multiple brain regions (Howard *et al*., 2000; De Peri *et al*., 2012). Hence, there is a need to refine the network incorporating the available spatio-temporal data. While the HiPPIP has led to a large theoretically possible interactome, the biological networks *in-vivo* are likely to be a subset of the computationally predicted network. This is mainly because the genes must be co-expressed and co-localized in-order to interact. In addition, the biological relavance of the experimental evaluations carried out in non-CNS tissues is debatable (Ganapathiraju *et al*., 2016). It would perhaps be more meaningful to evaluate the suspected targets in brains of patients with psychiatric illness.

Antipsychotics (AP) have been in use since 1950s (Shen *et al*., 1999). The first-generation APs were derived from a number of older drugs exploring antibiotic and anaesthetic effects, as well as drugs used in traditional medicine. At present, the commonly used drugs are second-generation APs, with their therapeutic effects largely being mediated by dopaminergic and serotonergic receptor blocking activities (Naheed *et al*., 2001). Antipsychotics have been associated with long-term side effects such as weight gain (Susilova *et al*., 2017), adverse metabolic effects, aggravating cognitive dysfunction (Zhang *et al*., 2017) and many others. Lithium and valproic acid have been administered to patients with bipolar disorder, but their mechanism of action is still incompletely understood (Rogers *et al*., 2017). There is thus a pressing need for new drugs in psychiatry.

Hence, by integrating data points from non-psychiatric ExAC, centenarian genomes, Allen Brain Atlas, expression profiles of psychiatrically ill post-mortem brain samples, SZ interactome and gene-drug interaction network, we present a multi-scale analysis to improve the current understanding of the genomic and pharmacological complexity of neuropsychiatric disorders (Girard *et al*., 2012; Ripke *et al*., 2014; Farrell MS *et al*., 2015; Ganapathiraju *et al*., 2016; Exome Aggregation Consortium *et al*., 2016; Lanz *et al*., 2015). The components of genomic variation associated with the disease, are likely to influence the disease phenotype through changes in protein biology. Our “multi-scale” analysis addresses the 4 mechanisms by which genomic variation could lead to the disease phenotype; by affecting the protein activity/function (identification of lethal non-synonymous variations), quantity of protein (in normal and post-mortem brain tissues), timing (multiple developmental stages) and location (multiple brain regions) of protein production. In the absence of quantitative protein expression data, gene expression (mRNA abundance) is taken as a surrogate of the protein levels. The translational control and protein degradation pathways could not be a part of the analysis.

To begin with, the variants present in SZ genes were mined from Ensembl. The variants that were absent in genomes of healthy centenarians and non-psychiatric ExAC database was identified and defined as *variants of criticality*. We harnessed the spatio-temporal gene expression data of SZ candidate genes from BrainSpan Atlas, and integrated them into the existing SZ interactome to identify critical genes and interactors as potential targets for therapeutic interventions. We hypothesize that the resultant dynamic network and the interactome would be a better approximation of the real biological network of SZ genes in a developing human brain. We harnessed the transcriptome data of psychiatrically ill postmortem brain tissues from Gene Expression Omnibus (GEO) (Lanz *et al*., 2015), to identify differentially expressed genes (DEGs) present in SZ interactome. Some of the interactors provided insights into psychiatric disorders and associated comorbidities like inflammation, immune dysfunction and visual deficits. The druggable DEGs and their pathways were identified, presenting a probable subset of targets for repurposing existing drugs for psychiatric disorders.

## MATERIALS AND METHODS

The analysis is represented as a graphical abstract (Figure 1)

**Figure 1:**
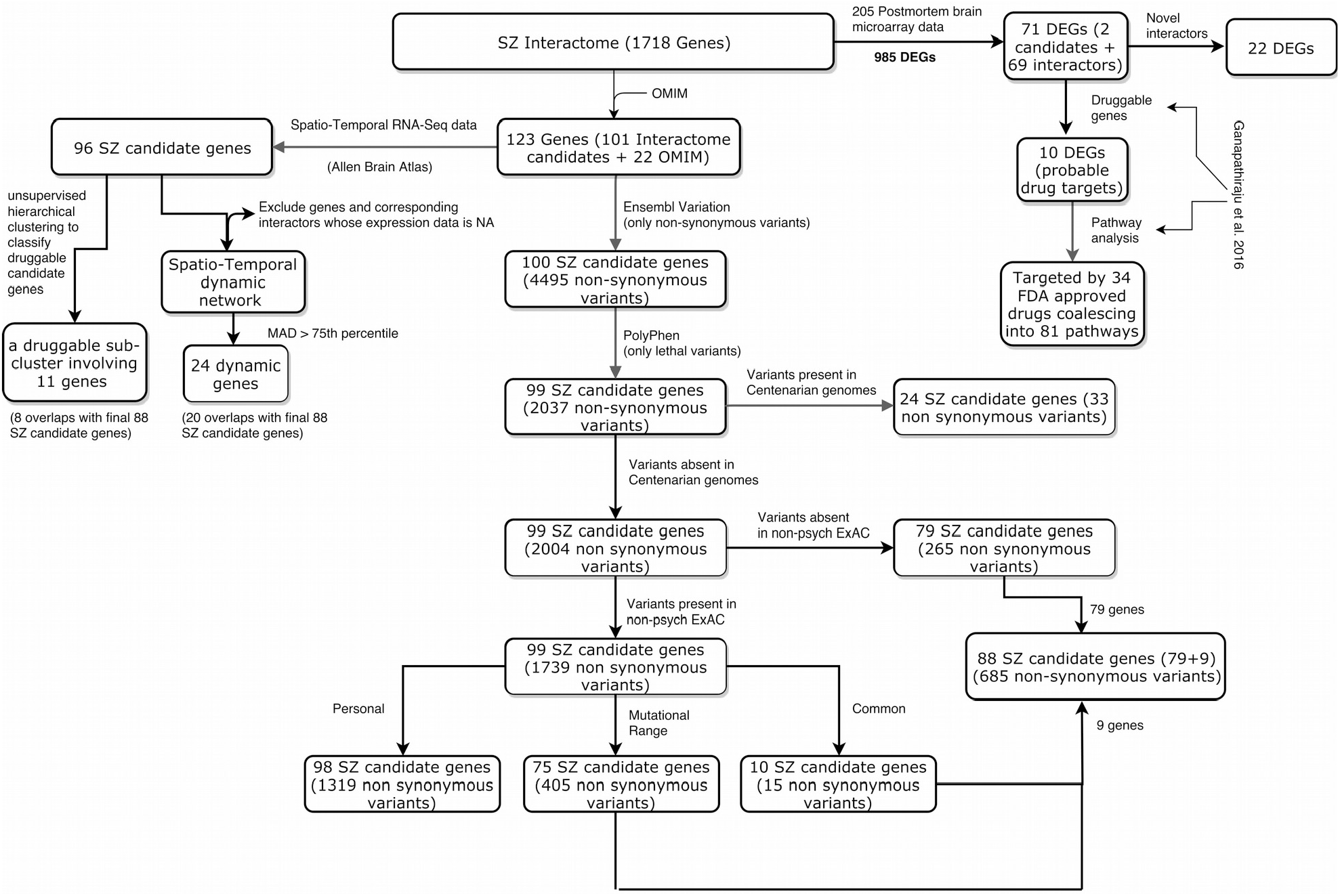
*Workflow: Multi-scale analysis of SZ genes*

### Database mining of single nucleotide variants present in the candidate genes

123 genes (101 interactome candidate genes + 22 OMIM genes) associated with SZ were retrieved from literature (Ganapthiraju *et al*., 2016). The 101 interactome candidates were themselves derived from 77 GWAS (Ripke *et al*., 2014) and 25 historic/pre-GWAS genes (Farrell MS *et al*., 2015), with *GRM3* being common. Apart from which, the 22 OMIM genes associated with SZ were also retrieved from the supplementary material of Ganapthiraju *et al*.,2016. Their genomic co-ordinates (GRCh37) were extracted from Ensembl’s Biomart (Yates *et al*., 2016). The gene symbols and their co-ordinates are given in Supplementary File 1. Well annotated non-synonymous variants were mined from Ensembl Variation (Yates *et al*., 2016) (GRCh37) for 123 candidate genes, of which 100 had well annotated non-synonymous variations. The functional consequences of the variants were predicted using Polymorphism Phenotyping v2 (PolyPhen-2) (Adzhubei *et al*., 2010). The *probably damaging* variants as predicted by PolyPhen were queried in (i) genomes of healthy centenarians (N=93) (http://clingen.igib.res.in/sage/) and (ii) non-psychiatric ExAC database (N=47,082) (Exome Aggregation Consortium *et al*. 2016). Non-psychiatric ExAC (version 0.3) variants with genotype quality≥20 and read depth≥10 were used for the above analysis. The genes and variants were then screened against several literature, databases, including *Online Mendelian Inheritance in Man* (OMIM) (Amberger *et al*., 2015) to check for association with other chronic illnesses apart from SZ.

### Construction of spatio-temporal dynamic network

The RNA-seq dataset from the BrainSpan Atlas of the developing human brain (Tebbenkamp *et al*., 2014) were retrieved for SZ candidate genes using the R package *ABAEnrichment* (Grote *et al*., 2016), which contains expression data only for protein-coding genes (aligned to GRCh37). To increase the power in detecting developmental effects by using highly overlapping brain regions, the dataset for the enrichment analysis was restricted to the 16 brain regions sampled in 5 developmental stages. Amongst 123 candidate genes, the spatio-temporal Reads Per Kilobase of transcript per Million mapped reads (RPKM) values were available only for 96 (89 interactome candidates + 7 OMIM candidates) genes, and the remaining 27 genes and their interacting partners, if any, were excluded from the analysis. The raw data was z-score normalized (gene-wise), and the median absolute deviation (MAD) (scaling factor, *k*=1) of the expression was calculated for each gene across all 16 tissues in 5 developmental stages. To facilitate the understanding of PPI network dynamics with respect to the brain regions as well as developmental stages, we developed an open source network visualisation toolkit. The toolkit is written in JavaScript using ReactJS [https://facebook.github.io/react], SigmaJS [http://sigmajs.org], and D3 [https://d3js.org] packages. This toolkit is accessible publicly on https://placet.noop.pw and the source code can be accessed from https://github.com/prashnts/placet. Hyperlink-Induced Topic Search (HITS) was used to rank the genes in the network based on the degree of the nodes. The normalized spatio-temporal gene expression data was integrated into the interactome, with the node sizes representing the expression levels of the corresponding genes. In order to classify the druggable genes from the remaining causative candidates, we employed hierarchical classification of spatio-temporal expression data of 96 SZ genes.

### Post-mortem microarray data analysis

Microarray expression profiles of 54675 Affymetrix probesets of PFC (Pre-frontal cortex), HPC (Hippocampus) and STR (Striatum) from 205 psychiatric subjects (in total) with SZ, BP, MDD and clinically matched controls were downloaded from GEO (ID: GSE53987) (Lanz *et al*., 2015). The downloaded data was MAS 5.0 normalized and log2 transformed to make sure that the data follows a Gaussian distribution. The distribution was looked for samples that might show variations in gene expression. The mean of expression of each gene across its corresponding samples was calculated. The fold change (FC) was calculated between the gene expression means of cases and corresponding controls. Student’s t-test was used to test for difference in gene expression between cases and controls. The False Discovery Rate (FDR) of t-test p-values was calculated for multiple hypothesis testing. A two fold change (FC>2) in gene expression in cases compared to controls along with a p-value<0.01, was considered to be differentially expressed. Annotation of Affymetrix probe IDs was perfomed using Affymetrix Netaffx Batch Query (http://www.affymetrix.com/analysis/index.affx). The union set of all DEGs in 9 different cases was identified. Ganapathiraju *et al*., had identified 504 novel PPI addition to the 1397 PPI, comprising of 1901 interactions in total. We have arrived at 1718 genes, by identifying the union set of all genes present in 1901 interactions. We then overlapped the union set of all DEGs identified, with all 1718 genes in the SZ interactome, for the downstream analysis.

### Identification of druggable genes and pathways

A 2-D matrix representing 286 biological pathways involving 122 druggable genes, was constructed from the literature (Ganapthiraju *et al*., 2016). The DEGs identified from the postmortem brain tissues were overlapped with 122 druggable genes, to identify the drug targets in the interactome that are differentially expressed from the post-mortem microarray study. An independent analysis was carried out using ConsensusPathDB (Release 32) (Kamburov *et al*., 2011) to identify more druggable genes (apart from 122) in biological pathways to which SZ candidate genes have been attributed (P<0.01).

### Statistical analysis and data visualization

All the statistical tests and data visualizations were performed using R, including MAS5.0 normalization and statistical corrections of microarray gene expression data.

## RESULTS

### 1. Functional consequences of non-synonymous variants

In order to characterise the functional implications of the non-synonymous variants in SZ candidate genes, we mined data from Ensembl Variation (EV) (Yates *et al*., 2015). Out of 123 SZ candidate genes (101 interactome candidate genes + 22 OMIM genes), EV reported 4495 well annotated non-synonymous variants in 100 SZ candidate genes for which the PolyPhen scores were retrieved (Supplementary Table 1A) (Adzhubei *et al*., 2010).

#### 1.A. Identification and shortlisting of lethal variants using genomes of healthy centenarians and non-psychiatric ExAC database

According to PolyPhen analysis, it was observed that 2037 (of 4495) variants were lethal *(probably damaging)* which mapped to 99 (of 100) SZ genes. In order to narrow down the number of deleterious variants, of the 2037 variants, we eliminated 33 variants belonging to 24 genes that were observed in genomes of healthy centenarians (N=93) (Supplementary Table 1B)(http://clingen.igib.res.in/sage/). Of the 2004 variants absent in centenarians (Supplementary Table 1C); i) we found that 265 variants (mapping to 79 genes) were also absent in non-psychiatric ExAC database and were defined as *variants of criticality* (Supplementary Table 1D) (Exome Aggregation Consortium *et al*., 2016). ii) we retained the remaining 1739 lethal variants i.e., those absent in centenarians but present in non-psychiatric ExAC, that mapped to 99 genes that could turn deleterious later on under certain circumstances (Supplementary Table 1E). These 1739 variants were further classified into three categories based on their allele frequencies (AF) (AF<0.0001: Personal; AF: 0.0001 to 0.01: Mutational range; AF>0.01: Common). Amongst the 1739 lethal variants, 1319 (mapping to 98 genes) were personal, 405 (mapping to 75 genes) were at mutational range and only 15 (mapping to 10 genes) were common in populations. We limited our analysis to common and mutational range variants but not the personal variants, since association of individual personal variants in complex disorders like SZ may represent a very small proportion of the possible risk factors, and unlikely to contribute susceptibility, at the population level. Hence, the 265 *variants of criticality* might act alone, or in combinations with the 15 common and 405 mutational range variants, i.e., 685 variants in total, mapping to 88 SZ genes (79 high risk genes + 9 genes unique to the mutational range and common variant genes), to contribute to the disease phenotype (Figure 1). Hence, we present a panel of potentially deleterious 685 variants, that could be further investigated for behavioral phenotypes and brain patho-biology in animal models of neuropsychiatric disorders (Supplementary Table 1F). It was also witnessed that, six (*CSMD1, CACNA1C, PLCH2, NRG1, ADAMTSL3* and *TCF20*) out of 88 SZ candidate genes, had a relatively higher burden of non-synonymous variants (>20 variants per gene) (Supplementary Figure 1). It is interesting to note that although the number of variants gets reduced at every step during the variant filtration process, the number of genes remain fairly the same. This could be because the risk variants are distributed among genes identified by the GWAS and other association studies, but seldom cluster on to a particular locus.

#### 1.B. Distribution of non-synonymous variants present in SZ genes in global populations

In order to gain an overall snapshot of the allele frequencies of the variants present in SZ genes in global populations, we queried all the original 4495 variants, in non-psychiatric ExAC database. On analysis, it was observed that 4045 variants were mapped to 99 genes, thereby discarding the 450 variants that were absent in the non-psychiatric ExAC database. The AFs of 4045 variants in 6 populations reported in the non-psychiatric ExAC database, was also retrieved (Supplementary Table 2). The analysis revealed that the number of variants observed in each of the population were directly proportional to their sample size. However, the proportion of the personal variants was higher (n=1843) in the out-bred European population (NFE) but was absent in the inbred Finnish (FIN) and East-Asian Tibeto-Burman (EAS) population (Figure 2). Although the personal variants were absent in FIN and EAS, the prevalence of psychiatric disorders was found to be as high in both the populations (Lehtinen *et al*., 1990). Thus, the absence of personal variants in FIN and EAS populations could be an artifact of the under-representation of the corresponding cohorts in the ExAC database.

**Figure 2:**
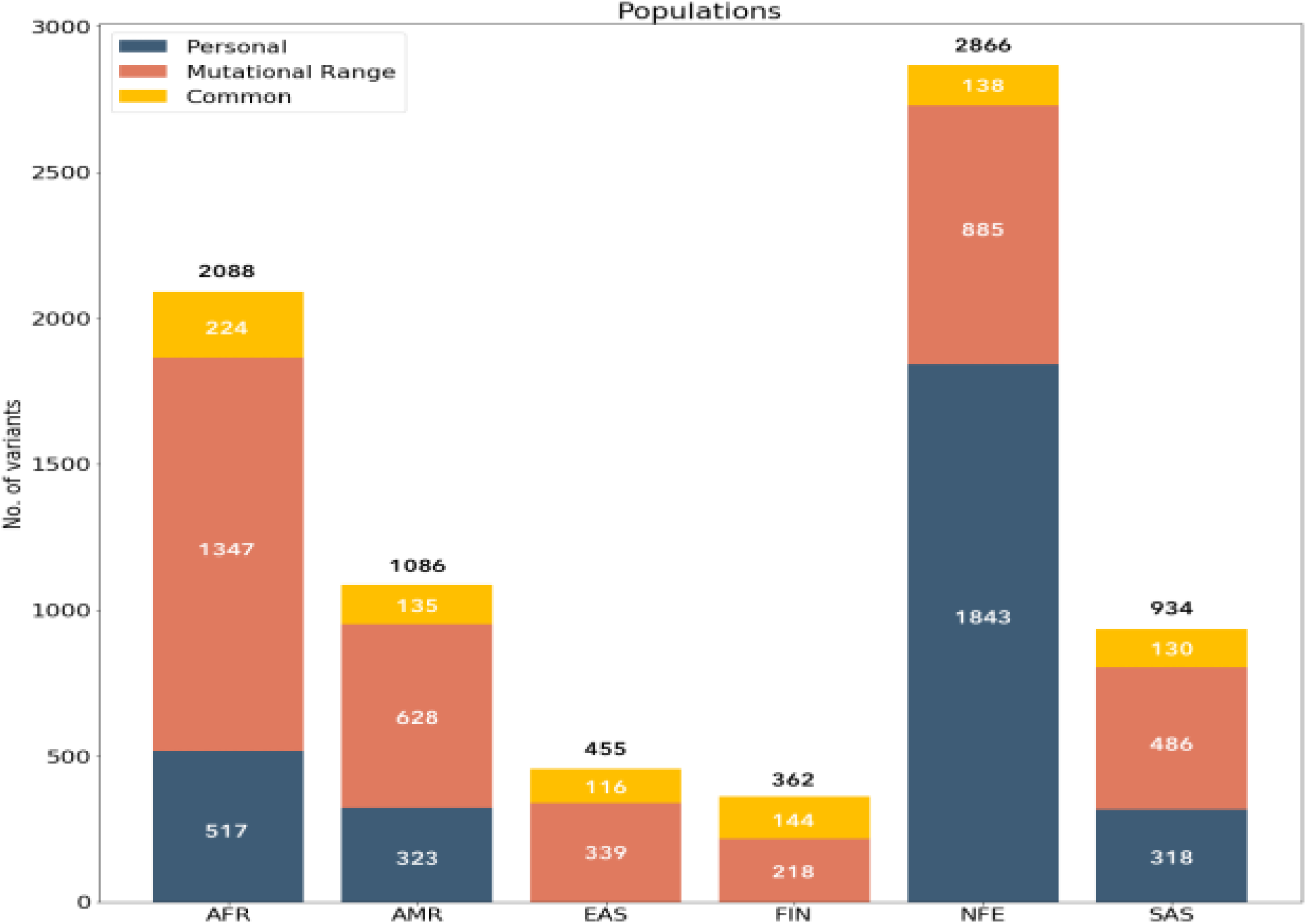
*Population distribution of personal, mutational range and common non-synonymous variants in SZ genes from non-psychiatric ExAC database. Amongst 4495 variants, 4045 mapping to 99 (out of 100 SZ genes) were identified in 6 populations. The variants observed in each population are directly proportional to their sample size. The bar diagram represents personal variants in blue color, the mutational range variants in red and the common variants in yellow color. AFR: African/African American; AMR: Latino; EAS: East Asian; FIN: Finnish; NFE: Non-Finnish European; SAS: South Asians*.

#### 1.C. Literature mining of variants present in SZ candidate genes that have been associated with multiple chronic illnesses

In order to verify the association of SZ candidate genes with other chronic illnesses, we utilized OMIM, literature in pubmed and other online sources, which revealed 94 disease associated non-synonymous variants present in 37 SZ genes, i.e., almost 40% of all schizophrenia candidates (Amberger *et al*., 2015). These variants were associated with other disorders include BP, MDD, Autism, Epilepsy, Seizure, Alzheimer’s, diabetes, hypertension, etc (Supplementary Table 3). Amongst 94 variants only 22 were predicted to be lethal by PolyPhen analysis which highlights the ambiguities inherent in the current methods in predicting the protein deleteriousness. Out of these 22 presumed lethal variants, twelve (rs2904552, rs3970559, rs34845648, rs8192466, rs45571736, rs2229961, rs34622148, rs1801500, rs769455, rs1801158, rs3970559 and rs2904552) were found to be absent in centenarian but present in the non-psychiatric ExAC database. However, none of the 22 were absent both in the centenarians and the non-psychiatric ExAC database.

### 2. Analysis of Spatio-Temporal Interactome

Although the PPI map for SZ (Ganapthiraju *et al*., 2016) presented all the possible interactions, a large proportion of the genes represented in the interactome are not coexpressed in a given location of the brain at a particular developmental stage. Therefore, we retrieved and integrated the spatio-temporal gene expression data from BrainSpan Atlas into the existing SZ interactome, thereby redefining the network as a function of space (16 brain regions) and time (5 developmental stages) (http://placet.noop.pw/) (Supplementary Table 4; Supplementary Figure 2) (Tebbenkamp *et al*., 2014; Grote *et al*., 2016).

#### 2.A. Extent of difference in gene expression between hub genes and non-hub genes

Hyperlink-Induced Topic Search (HITS) was used to rank the genes as hubs (top 10 genes) or non-hubs (bottom 10 genes). With this study, we hypothesized that highly connected genes i.e., the hub genes, must be expressed significantly higher compared to non-hub genes inorder to interact with larger set of proteins. However, no difference was found between the gene expression means of hub genes and non-hub genes.

#### 2.B. Characterization of gene expression dynamics in regions of adolescent and adult brain

In order to identify the genes that exhibit high variations in expression pattern in a normal human brain, we carried out median absolute deviation (MAD) score analysis for the 96 SZ candidate genes across the spatio-temporal gene expression data. The analysis revealed that the expression of 24 genes was highly dynamic across space and time amongst which 13 (*RGS4, HTR2A, APOL2, GRIN2A, CNNM2, CACNA1C, ZDHHC8, HCN1, DPYD, OTUD7B, ZNF536, C3orf49* and *CLCN3)* were highly expressed in adult and/or adolescent brain tissues compared to child, infant and prenatal brain tissues (Supplementary Table 5).

#### 2.C. Expression based similarity search for druggable SZ candidate genes

Despite large GWAS findings, it still remains unclear which, if any, of the newly identified GWAS loci will serve as good starting points for drug development in SZ (Dolgin *et al*., 2014). Most drug targets may not be ubiquitously expressed but enriched and localized in distinct tissues relevant to the disorders, even under normal conditions (Kumar *et al*., 2016). Hence, there is a need to classify the drug targets from candidate genes based on their expression patterns. In order to classify the druggable genes from the remaining causative candidates we employed hierarchical classification of spatio-temporal expression data of 96 SZ candidate genes and identified a sub-cluster of 11 genes (*IGSF9B, NAB2, DAO, CYP26B1, PLCH2, CHRNA3, SLC6A3, DRD2, DRD3, DAOA* and *TAAR6*), which were enriched only in certain regions of the brain at certain developmental stages (Figure 3).

**Figure 3:**
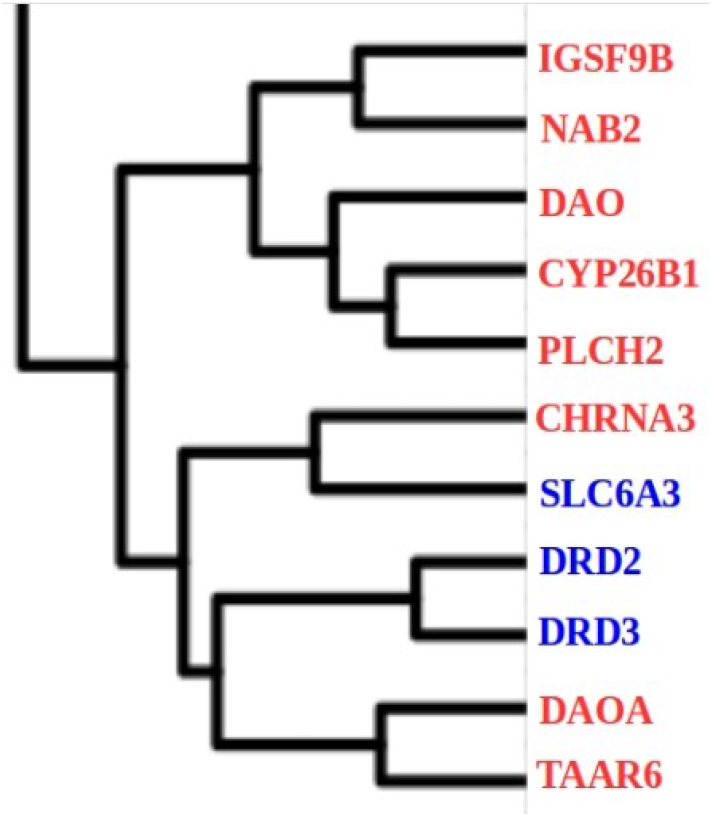
*11 druggable candidate genes classified based on similarity in gene expression signatures. Blue: known drug targets for SZ; Red: putative drug targets for SZ. Amongst which SLC6A3, DRD2, DRD3, DAO, DAOA and CHRNA3 involve in neurotransmission*.

Amongst the sub-cluster of 11 genes, three (*SLC6A3, DRD2* and *DRD3*) have been well-established drug targets for SZ (Figure 4). *SLC6A3* was found to be active in child and adolescent thalamus, a region which has been implicated in SZ (Figure 4C) (Alelú-Paz *et al*., 2008; Pergola *et al*., 2015). *DRD2* (Figure 4A) and *DRD3* (Figure 4B) were found to be active only in striatum across all developmental stages. Striatum has been associated with the pathophysiology of SZ, BP, ASD and adolescent depression (Brunelin *et al*., 2016; Simpson *et al*., 2010; Fuccillo *et al*., 2016; Caseras *et al*., 2013; Gabbay *et al*., 2013). *DAO* and *DAOA* which also belong to the sub-cluster, have been receiving attention as potential alternative therapeutic means to enhance *NMDAR* function in SZ patients (Verrall *et al*., 2010; Sehgal *et al*., 2015). DAO was found to be enriched in anterior cingulate cortex (MFC) which has also been implicated in SZ (Pomarol-Clotet *et al*., 2010). Interactors of *NAB2* are targeted by nervous system drugs in the management of epilepsy, which often co-occurs with SZ (Ganapthiraju *et al*., 2016; Cascella *et al*., 2009). NAB2 is highly expressed in cerebellar cortex, a brain region that shows modest association with SZ endophenotypes (Kim *et al*.,2014; Bang *et al*., 2018; Andreasen *et al*., 2011). Trace amine-associated receptors (*TAAR*) are GPCRs that are often activated after the blockade of dopaminergic receptors by antipsychotics (Kleinau *et al*., 2011). It is also interesting to note the silence of *TAAR6* in most regions of the brain which are of interest in SZ, except in the MFC (Child). Whether *TAAR* activation observed is a pathobiological correlate of the disorder, or a consequence of receptor blockade by antipsychotics, and thus a psycho pharmacologically mediated process, needs to be understood. Although the expression of the remaining 4 genes (*IGSF9B, CYP26B1, PLCH2* and *CHRNA3*) in the sub-cluster show druggability signatures, no strong evidence was available from existing literature. The spatio-temporal expression maps of 96 SZ genes are available in Supplementary File 2. Multiple evidences indicate that the therapeutic outcome of antipsychotics is mediated not just by the classical dopamine D2/D3 receptors, but also other targets including the D1 receptors. Moreover, the clinical outcome i.e., full or partial recovery from the first incidence of psychotic episodes may take weeks or even months, although the relevant receptors are blocked within hours (Miller *et al*., 2009). This multitude of effects, and side effects, and the latency of clinical outcome suggests a complex interplay of multiple biological pathways. These processes needs to be understood using post-mortem gene expression data, as well as data from animal model systems, to able to identify potential alternative drug targets for psychiatric disorders.

**Figure 4:**
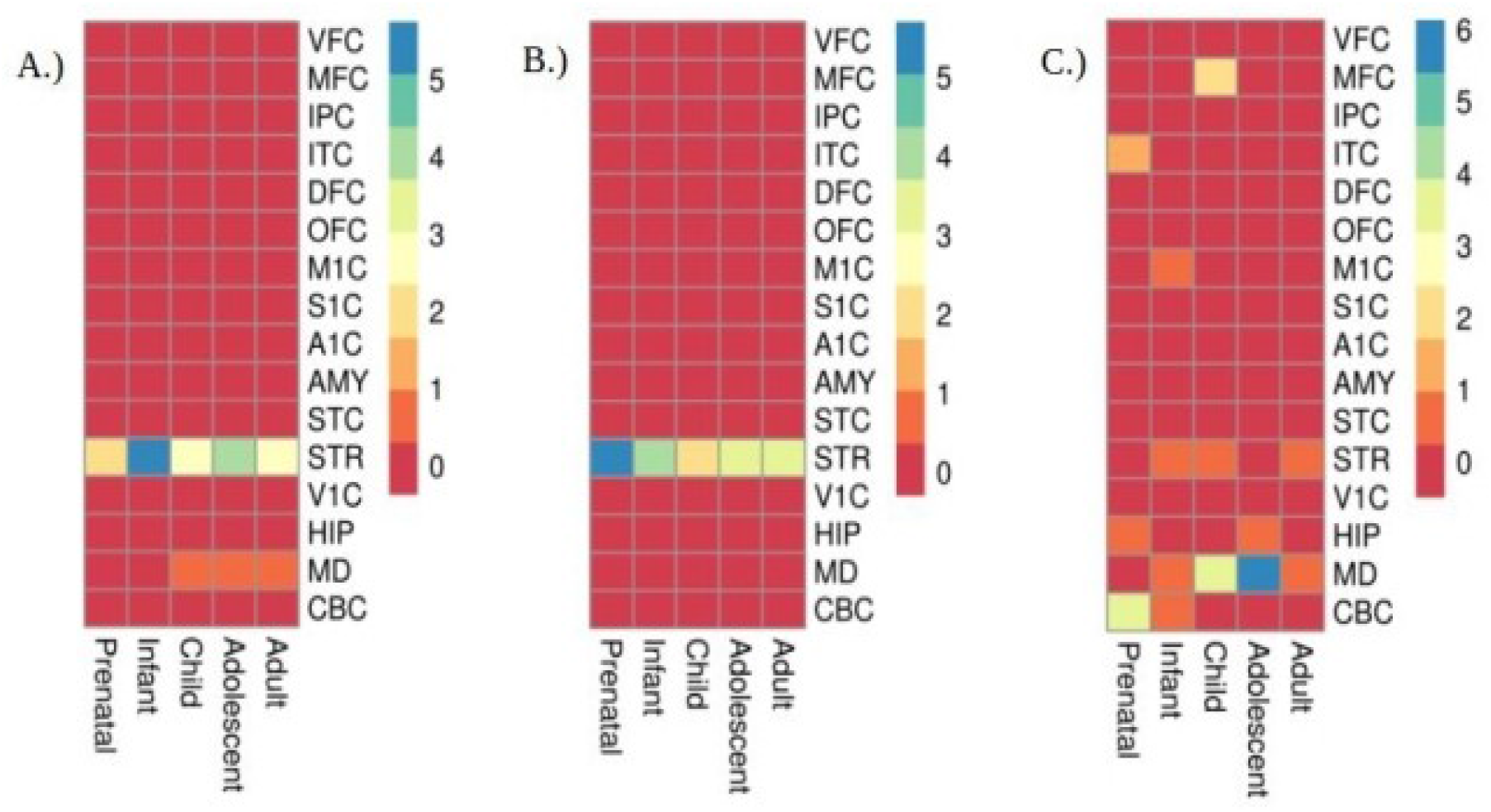
*Spatio-temporal expression profiles (Z_score RPKM) of druggable SZ candidate genes A.) DRD2, B.) DRD3 and C.) SLC6A3 in a developing human brain*.

#### 2.D. Analysis of differentially expressed genes in psychiatrically ill post-mortem brain tissues and their overlap with SZ interactome

In order to identify the genes present in the SZ interactome that are dysregulated in psychiatric patients, the microarray expression profiles from 205 post-mortem brains of patients were examined (Lanz *et al*., 2015). Analysis of expression profiles of PFC, HPC and STR revealed 985 unique DEGs (FC>2; *P*<0.01) from nine different conditions (Supplementary File 3). The raw t-test *P*-values were used since FDR corrected *P*-values were not significant for most genes to be called as differentially expressed. In order to validate the role of genes from the SZ interactome (Ganapthiraju *et al*., 2016) in post-mortem brain tissues, we overlapped the gene IDs of 985 DEGs and 1718 genes in SZ interactome. We obtained an overlap of 71 genes (2 candidate genes + 69 interactors) that were present in the SZ interactome and also differentially expressed in post-mortem brain tissues; of which 22 being novel interactors as predicted by Ganapthiraju *et al*., 2016. Fourteen of the 22 dysregulated genes in our analysis revealed direct or indirect relationships with neuropsychiatric disorders and co-morbidities from previous studies (Table 1). The remaining 8 novel interactors (*MYOZ2, CARS, GSC2, MKI67, ZC3H15, HOPX, CDC42S1E* and *VANGL1*) though dysregulated in our analysis, had no previous mention in the literature with psychoses. On analysis, it was observed that only two SZ candidate genes (*SLC6A4* and *CACNB2*) were differentially expressed in HPC (BP) (log2FC=1.31; P=0.005) and PFC (MDD) (log2FC=-1.11; P=0.008) respectively (Supplementary File 3). Interestingly, 14 out of 71 DEGs were previously identified as druggable targets of various FDA approved drugs in the gene-drug interactome study by Ganapthiraju *et al*., 2016. The above analysis is illustrated in Supplementary Figure 3.

**Table 1:**
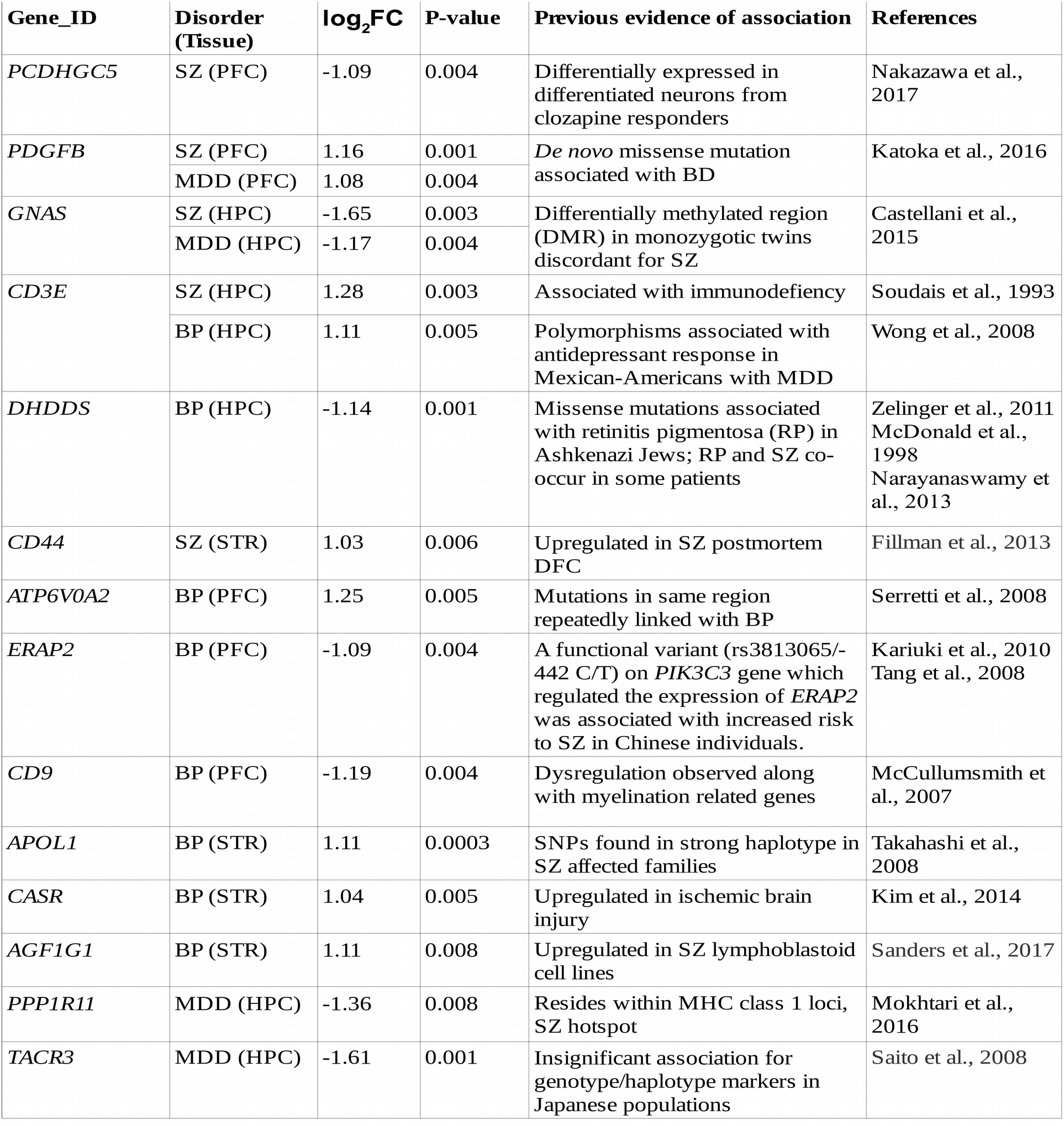
*Dysregulated novel interactors identified from post-mortem microarray analysis.*

### 3. Analysis of druggable genes and pathways

A 2-D matrix representing 286 biological pathways involving 122 druggable genes was reconstructed from the literature (Ganapthiraju *et al*., 2016). From the above post mortem gene expression data from patients, we identified 14 druggable DEGs. Of these 14 DEGs, four (*PTGS1, ERBB2, PTGER3* and *ESR2*) were found to be downregulated in at least one of the nine conditions. Since, drug induced inhibition of highly expressed would be easier (*It’s all druggable*., 2017), the four downregulated targets were excluded from downstream analysis. This resulted in 10 upregulated and probable drug targets including *ACE, CD44, mTOR, RARA, PTPN1, LDLR, CD3E, NOS3, CFTR* and *CASR,* targeted by 34 FDA approved drugs (Table 2) (Supplementary File 4) belonging to 81 biological pathways (Supplementary Table 6).

**Table 2:** *Shortlisted druggable genes and their corresponding FDA approved molecules*.

| Drug Target<br>(Gene symbol)<br>(n=10) | Gene name | Interacting drugs (FDA approved) | No. of FDA approved drugs<br>(n=34) | Relevance to psychoses/brain biology | Support for druggability (animal models, etc) |
| --- | --- | --- | --- | --- | --- |
| *ACE* | Angiotensin converting enzyme | Ramipril, Fosinopril, Trandolapril, Benazepril, Enalapril, Moexipril, Perindopril, Quinapril, Rescinnamine, Captopril, Cilazapril, Spirapril and Temocapril | **13** | Nadalin et al., 2017<br>Gadelha et al., 2015<br>Gadelha et al., 2015<br>Eckman et al., 2006 | DK.Hobgood. 2013<br>AbdAlla S et al. 2013<br>Gadelha A et al. 2015 |
| *CD44* | Cell-surface glycoprotein | Hyaluronan | **1** | Ponta et al., 2003 | NA |
| *mTOR* | Serine/threonine-protein kinase | Everolimus, Temsirolimus, Sirolimus and Pimecrolimus | **4** | Kim et al., 2002<br>Pham et al., 2016<br>Wang et al., 2017 | Tufts SZ mTOR studies<br>Lipton ans Sahin. 2014.<br>Zhou et al. 2013 |
| *RARA* | Retinoic acid receptor alpha | Acitretin, Adapalene, Tazarotene, Alitretinoin and Etretnate | **5** | Phillippe et al., 2001<br>Haybaeck et al., 2015<br>Lerner et al., 2016 | Malaspina 2008.<br>Jarvis et al., 2010<br>Haybaeck et al. 2015<br>Rioux 2005 |
| *PTPN1* | Tyrosine-protein phosphatase non-receptor type 1 | Tiludronate | **1** | Imming et al., 2006<br>Freedman et al., 2001<br>Yin et al., 2017 | RJ He et al. 2014<br>Carty NC et al., 2012 |
| *LDLR* | Low-density lipoprotein receptor | Porfimer | **1** | Nourooz-Zadeh et al., 1996<br>Gibbons et al., 2010 | Gibbons et al., 2010 |
| *NOS3* | Nitric oxide synthase 3 | Miconazole, Tetrahydrobiopterin and L-arginine | **3** | Marsden et al., 1993<br>Pilar et al., 2008<br>Shinkai et al., 2002 | Pitsikas N. 2016<br>Lafioniatitis et al., 2016<br>Wass C et al., 2008 |
| *CFTR* | Cystic fibrosis transmembrane conductance regulator | Glyburide, Ivacaftor, Ibuprofen, Bumetanide | **4** | Gillen et al., 2012 | Puljak Livia. 2006 |
| *CASR* | Calcium-sensing receptor | Cinacalcet | **1** | Hendy et al., 2013<br>Gupta et al., 2007 | BA Twgit et al. 2013<br>Kim JY et al., 2014<br>Dal Pra I et al., 2015 |
| *CD3E* | T-cell surface glycoprotein CD3 epsilon chain | Muromonab | **1** | NA | FG Hoosian et al., 2015 |

#### 3.A. Investigation into druggable genes present in pathways belonging to SZ candidate genes

Amongst 71 DEGs identified from post-mortem brains, only 2 were candidate genes (*SLC6A4* and *CACNB2*), and 10 were druggable interactors. In order to identify more druggable genes upstream or downstream in the biological pathways associated with SZ, we used ConsensusPathDB (CPDB) (Kamburov *et al*., 2011) to identify pathways to which the original list of 123 SZ candidate genes belong (***P***<0.01). The CPDB analysis revealed overrepresentation of 46 (out of 123) genes in 54 biological pathways which includes Dopaminergic signaling, MAPK signaling, cAMP signaling, Axon guidance, Calcium signaling, Gα signaling, Celecoxib pharmacodynamics, T-cell receptor signaling, Alzheimer’s disease pathway, etc (Supplementary Table 7). Of these 54, only 3 pathways had putative drug targets (n=4 i.e., *COX-2, GNAQ, PLCB1* and *PDE10A*) and their spatio-temporal expression (Z-score_RPKM) profiles are also provided (Supplementary Figure 4,5,6,7).

##### 3.A.1 Celecoxib pharmacodynamics pathway

Celecoxib, a *COX-2* inhibitor is an anti-inflammatory drug used to treat osteoarthritis, RA, Ankylosing spondylitis (AS), acute pain in adults and juvenile arthritis. It has also been investigated as an adjuvant in several psychiatric disorders like MDD, BP and SZ (Fan *et al*.,2013; Muller *et al*., 2008; Na *et al*., 2014; Rosenblat *et al*., 2014; Fond *et al*., 2014). On analysis, the SZ genes, *CACNB2, PTGIS, ATP2A2* and *AKT1* were found to be involved in Celecoxib pharmacodynamics. Although the expression of the above over-represented genes has been characterized in previous analysis, the spatio-temporal enrichment of *COX-2* with respect to celecoxib metabolism has never been reported. Spatio-temporal expression profiles of *COX-2* were retrieved from BrainSpan atlas database using ABAEnrichment. Further analysis revealed that *COX-2* levels were high in all the 11 cortical regions of Infant, Child and Adolescent human brain, especially in Infant (S1C) and Adolescent (A1C) (Supplementary Figure 4).

##### 3.A.2 Gα signaling pathway

The associations of G-protein coupled receptors and Gα subunits in phosphoinositide signaling with psychoses has been well established (Catapano *et al*., 2007; Rajy *et al*., 1997). The two genes (*PLCB1* and *GNAQ*) were found to be involved in Gα signaling pathway and deletions in the former were observed in SZ patients (Lo *et al*., 2013). The analysis revealed that the expression of *GNAQ* was high in Prenatal (CBC) (Supplementary Figure 5). However, expression of *PLCB1* was found to be high in V1C (Adult, Child and Infant) and Infant (STR) (Supplementary Figure 6).

##### 3.A.3 cGMP-PKG signaling pathway

Phosphodiesterase 10A (*PDE10A*) is a basal ganglia specific hydrolase that regulates cAMP/cGMP signaling cascades. Animal studies have revealed that *PDE10A* inhibitors could provide efficacy on the positive and negative symptoms of SZ, and these are currently being evaluated in clinical trials (Kehler *et al*., 2011). However, their spatio-temporal expression patterns were unknown. Our analysis using BrainSpan data revealed that *PDE10A* was enriched only in STR, which is a part of basal ganglia. Maximum expression of *PDE10A* was observed in infant (STR) (Supplementary Figure 7).

## DISCUSSION

Despite extensive genetic studies of neuropsychiatric disorders, the molecular mechanisms of patho-biology are still unknown. A computational systems biology study had identified protein interactions of SZ candidate genes and predicted a large number of novel interactions and interactors amongst which several were targets of FDA approved drugs (Ganapthiraju *et al*., 2016). With the availability of the genomes of healthy centenarians, non-psychiatric ExAC and BrainSpan data, we have made an attempt to identify relevant candidate genes in the network that could influence the risk of psychoses. By leveraging centenerian genomes and non-psychiatric ExAC data, the risk was narrowed down to 685 variants, spread over 88 SZ candidate genes, that could be investigated further using animal models. Literature mining suggested that ~40% of all SZ GWAS genes have shared genetic risk for one or more chronic illnesses, which needs to be validated by meta-analysis of genetic and clinical phenotype data. BrainSpan data suggested 13 dynamic and highly expressed genes in adult and adolescent brain regions, which might play a crucial role in the onset of psychiatric illnesses. Expression based similarity search of druggability in normal human brain, suggested the prioritization of 11 SZ candidate genes that could be potential targets of novel or repurposed drugs. Further, in order to identify the genes present in the SZ interactome that are dysregulated in psychiatric patients, the microarray expression profiles from 205 post-mortem brains were looked into, and the DEGs were overlapped with the union set of all genes present in the SZ interactome. Twenty two novel interactors present in the SZ interactome were found to be dysregulated in post-mortem brains. These proteins previously had null or minimal associations with psychoses, thereby now validating a subset of the novel interactors as predicted by Ganapthiraju *et al*., 2016. We also observe the dysregulation of *DHDDS*, a gene that has been strongly associated with Retinitis Pigmentosa, which occasionally co-occurs in certain schizophrenia cases (Table 1) (McDonald *et al*., 1998). Although no direct evidence for psychoses was found for 8 novel interactors that were dysregulated in psychiatric postmortem brains, some of them (*MYOZ2, GSC2, MKI67* and *VANGL1*) were discernable and need further investigation, as they point to critical processes. *MYOZ2* belongs to a family of sarcomeric proteins that bind to calcineurin, a phosphatase involved in calcium and calcineurin signaling, which are critical for SZ biology (Lidow *et al*., 2003; Miyakawa *et al*.,2003). *GSC2*, a homeodomain containing gene resides on 22q11, which is a hotspot for psychoses (Saleem Q *et al*., 2001). *MKI67* encodes a nuclear protein that is associated with cellular proliferation, and it has often been suggested that SZ is a disorder of inappropriate neuronal proliferation and pruning (Keshavan MS *et al*., 1994). Mutations in *VANGL1* are associated with neural-tube defects (Kibar *et al*., 2009) which have also been associated with increased risk in SZ patients (Zammit *et al*., 2007).

Of the 10 druggable interactors that are shortlisted for repurposition (Table 2), it would be meaningful to investigate the action of drugs in the context of receptor based (*CD44, RARA, LDLR, CASR* and *CD3E*) and non-receptor targets (*ACE, mTOR, PTPN1, NOS3* and *CFTR*), in ameliorating the whole spectrum of symptoms of SZ and other psychoses. The druggable genes that were further identified in pathways involving the SZ candidates, including *COX2, PLCB1, GNAQ* and *PDE10A*, were found to be highly expressed in the developmental stages that are pertinent to the onset of psychiatric illness. Thus, investments must be made into experimental validation in confirming the role of the above four genes and interacting small molecules, in ameliorating SZ like symptoms in animal models. The biological pathways, though diverse, cover a broader spectrum of cellular functions such as viability, proliferation and regulation of cell motility, which are generic, but may be critical to the pathobiology of schizophrenia. It is now fairly evident that the drugs that rely predominantly on modifying dopaminergic or seretonergic neurotransmission, may be inadequate to address the complexity of the biological processes, that we are now beginning to understand. One size, indeed, may not fit for all. Hence, there is a pressing need for adjunctive therapeutic strategies targeting the genes and pathways that are being detected by current research. Validation of these proposed drugs, drug targets and pathways in animal models and induced pluripotent stem cells (iPSC) derived neuronal lineages of SZ patients (Viswanath *et al*., 2018), could be useful to help unravel the biology of mental illness, and also accelerating the drug repurposing pipelines.

### Availability of data and materials

This spatio-temporal dynamic network is accessible publicly on https://placet.noop.pw and the Source code can be accessed from https://github.com/prashnts/placet. All PolyPhen annotated variants, DEGs identified from post-mortem microarray, lists of biological pathways etc., are available as Supplementary Files.

## AUTHOR DISCLOSURE

### Contributors

SKB conceptualized and designed the project. ACS and AKJ performed the gene expression data analysis. AKP performed the non-psychiatric ExAC variation analysis and PS constructed the spatio-temporal network. ACS and SKB wrote the manuscript. SJ provided intellectual support in interpreting the results and editing the manuscript.

## Acknowledgments

SKB is a recipient of the J.C. Bose National Fellowship. ACS thanks Mohandas Pai foundation for providing fellowship support through Centre for Open Innovation, IndianCST. We thank Raja Seevan, Sri Kumar and the IndianCST team for the infrastructure support. We thank NIMHANS for providing institutional support to SJ. We thank N. Balakrishnan for providing access to the computational facility at the Supercomputer Education and Research Centre, Indian Institute of Science. We also thank Vinod Scaria for providing access to the allele frequencies from his unpublished centenarian genome data and Beena Pillai for inputs on gene expression data analysis. We finally thank Meera Purushottam, Biju Viswanath and Ravi Kumar Nadella for critical reading of the manuscript.

